# Isolation of a *Bacillus safensis* from mine tailings in Peru, genomic characterization and characterization of its cyanide-degrading enzyme CynD

**DOI:** 10.1101/2021.11.27.470173

**Authors:** Santiago Justo Arevalo, Daniela Zapata Sifuentes, Andrea Cuba Portocarrero, Michella Brescia Reategui, Claudia Monge Pimentel, Layla Farage Martins, Paulo Marques Pierry, Carlos Morais Piroupo, Alcides Guerra Santa Cruz, Mauro Quiñones Aguilar, Chuck Shaker Farah, João Carlos Setubal, Aline Maria da Silva

## Abstract

Cyanide is widely used in industry as a potent lixiviant due to its capacity to tightly bind metals. This property imparts cyanide enormous toxicity to all known organisms. Thus, industries that utilize this compound must reduce its concentration in recycled or waste waters. Physical, chemical, and biological treatments have been used for cyanide remediation; however, none of them meet all the desired characteristics: efficiency, low cost and low environmental impact. A better understanding of metabolic pathways and biochemistry of enzymes involved in cyanide degradation is a necessary step to improve cyanide bioremediation efficacy to satisfy the industry requirements. Here, we used several approaches to explore this topic. We have isolated three cyanide-degrading *Bacillus* strains from water in contact with mine tailings from Lima, Peru, and classified them as *Bacillus safensis* PER-URP-08, *Bacillus licheniformis* PER-URP-12, and *Bacillus subtilis* PER-URP-17 based on 16S rRNA gene sequencing and core genome analyses. Additionally, core genome analyses of 132 publicly available genomes of *Bacillus pumilus* group including *B. safensis* and *B. altitudinis* allowed us to reclassify some strains and identify two strains that did not match with any known species of the *Bacillus pumilus* group. We searched for possible routes of cyanide-degradation in the genomes of these three strains and identified putative *B. licheniformis* PER-URP-12 and *B. subtilis* PER-URP-17 rhodaneses and *B. safensis* PER-URP-08 cyanide dihydratase (CynD) sequences possibly involved cyanide degradation. We identified characteristic C-terminal residues that differentiate CynD from *B. pumilus* and *B. safensis*, and showed that, differently from CynD from *B. pumilus* C1, recombinant CynD from the *Bacillus safensis* PER-URP-08 strain remains active up to pH 9 and presents a distinct oligomerization pattern at pH 8 and 9. Moreover, transcripts of *B. safensis* PER-URP-08 CynD (CynD_PER-URP-08_) are strongly induced in the presence of cyanide. Our results warrant further investigation of *B. safensis* PER-URP-08 and CynD_PER-URP-08_ as potential tools for cyanide-bioremediation.

## INTRODUCTION

Cyanide is a highly toxic compound used in several industrial processes (Mudder et al., 2004) given its capacity to form tight complexes with different metals (Dash et al., 2009) (Hendry-Hofer et al., 2019; Leavesley et al., 2008). Industries that generate cyanide-containing wastes must reduce its concentration before discarding them to the environment, and as such proper strategies have to be implemented for cyanide remediation (Kuyucak & Akcil, 2013). Cyanide bioremediation by bacteria that express nitrilases is one possible low-cost and environmental-friendly approach (Dash et al., 2009).

Nitrilases are a superfamily of proteins characterized by a tertiary structure consisting of an alpha-beta-beta-alpha fold and a dimer as a basic unit. This superfamily has been divided into thirteen branches with branch one corresponding to enzymes that cleave the nitrile group into ammonia and its respective carboxylic acid. The other twelve branches are structurally similar but their catalytic activity does not involve cleavage of nitriles (Pace & Brenner, 2001).

Two types of nitrilases can degrade cyanide through a hydrolytic pathway: cyanide hydratases (CHTs) and cyanide dihydratases (CynDs). CHTs convert cyanide into formamide using one water molecule in the reaction and are present in fungal genomes. The first enzyme described with this activity was from *Stemphylium loti* (Fry & Millar, 1972). Subsequently, CHTs from other fungal species have been studied such as: *Fusarium solani, Fusarium oxysporum, Gloeocercospora sorghi, Leptosphaeria maculans*, and *Aspergillus niger* (Akinpelu et al., 2018; Dumestre et al., 1997; Ping Wang; Hans D. VanEtten, 1992; Rinágelová et al., 2014; Sexton & Howlett, 2000). On the other hand, only the CynDs from *B. pumilus*, *P. stutzeri* and *Alcaligenes xylosoxidans* (Ingvorsen et al., 1991; Meyers et al., 1993; Watanabe et al., 1998) have been experimentally tested. The reaction catalyzed by CynDs generates formic acid and ammonia using two molecules of water. Both, CynDs and CHTs, typically form large helical aggregates of several subunits (Thuku et al., 2009). For example, the CynD of *Bacillus pumilus* is reported to form an 18-subunit oligomer (Jandhyala et al., 2003) whereas the homolog in *Pseudomonas stutzeri* forms a 14-subunit oligomer (Sewell et al., 2003)

Several *Bacillus* species have been shown to be capable of metabolizing cyanide using different routes, for instance: B-cyanoalanine synthase in *Bacillus megaterium* (Castric & Strobel, 1969), gamma-cyano-alpha-aminobutyric acid synthase in *B. stearothermophilus* (Omura et al., 2003), rhodanase in *Bacillus cereus* (Itakorode et al., 2019), and CynD in *Bacillus pumilus* (Meyers et al., 1993). On the other hand, some other cyanide-degrading *Bacillus* have still unknown metabolic routes (Al-Badri et al., 2020; Javaheri Safa et al., 2017; Mekuto et al., 2014).

The *Bacillus pumilus* group consists mainly of three species: *Bacillus altitudinis*, *Bacillus safensis* and *Bacillus pumilus*. These three species share more than 99 % sequence identity in their 16S rRNA gene (Liu et al., 2013), hampering taxonomical classification based solely on this locus. Studies using multiple phylogenetic markers have demonstrated that ~50 % of the *Bacillus pumilus* group genomes deposited in NCBI database could be misclassified (Espariz et al., 2016).

It is plausible to speculate that CynDs isolated from *Bacillus* strains from diverse environments could present different properties that could be better for industrial applications. Therefore, the characterization of CynD from other species can expand our understanding on the functioning and plasticity of this enzyme. Furthermore, some aspects of the biology of this enzyme have not been thoroughly studied. For instance, it is known that the oligomeric state of CynD is strongly pH-dependent (Jandhyala et al., 2003); however, the effect on oligomerization at pHs greater than 9 has not been reported. Also, it is unknown whether CynD is constitutively expressed in basal metabolism or is part of a specific physiological response, for instance, induced by the presence of cyanide.

Here, we describe the isolation of three indigenous *Bacillus* strains from mine tailing in Peru and their respective genome sequences. We selected a strain that was most efficient in cyanide degradation and investigated its phylogenetic relationship with other species of the *Bacillus pumilus* group. We identified a gene coding for a cyanide dihydratase (CynD) that is most likely the enzyme responsible for cyanide degradation in this selected strain. A recombinant CynD was expressed and purified, its catalytic parameters were determined, and the quaternary structure was studied at different pHs. We also demonstrated that CynD transcripts are strongly induced in the presence of cyanide.

## MATERIAL AND METHODS

### Isolation of cyanide-degrading strains

Water in contact with mine tailing was collected from a river near Casapalca and La Oroya mines located in San Mateo de Huanchor (Latitude −11.4067 Longitude −76.3361 at 4221 MASL). The sample was collected in 2 L sterile bottles and transported at 4 °C.

One hundred mL of the sample was added to an Erlenmeyer flask containing 20 ml of 21 g/L sodium carbonate, 9 g/L sodium bicarbonate, 5 g/L sodium chloride and 0.5 g/L potassium nitrate. Cultures were incubated for 12 h at 37 °C and after this time 1 mg/L final concentration of cyanide in the form of sodium cyanide was added. The cultures were incubated for another 24 h at 37 °C. Samples of the medium were streaked in petri dishes with nutrient agar (5 g/L peptone, 5 g/L yeast extract, 5 g/L sodium chloride and 1 % agar) and incubated at 37 °C for 24 h. Single colonies were isolated in nutrient broth (5 g/L peptone, 5 g/L yeast extract, 5 g/L sodium chloride) supplemented with 20 % glycerol and stored at −80 °C.

Strains stored at −80 °C were reactivated at 37 °C in nutrient agar by streaking a sample. One colony was inoculated in fresh nutrient broth and incubated at 37 °C overnight at 100 g. Next, the optical density at 600 nm (OD_600nm_) of the culture was adjusted to 0.8 and 1 mL was centrifuged at 6000 g for 3 min. The pellet was washed twice with 0.2 M Tris-HCl pH 8 and resuspended in 1 mL of 0.2 M Tris-HCl pH 8 supplemented with 0.2 M NaCN. After 2 h of incubation at 30 °C, the culture was centrifuged at 6000 g and 10 μL of the supernatant was taken and diluted in 90 μL of milliQ water. Then, 200 uL of 0.5 % picric acid in 0.25 M sodium carbonate was added and heated for 6 min at 100 °C (Williams & Edwards, 1980). Finally, absorbance at 520 nm was measured and compared to a standard curve of NaCN.

### Strain identification by 16S rRNA gene sequencing

To determine the bacterial genera and/or species of the isolated strains, we used a fresh culture in nutrient agar. Five colonies from these cultures were transferred to 50 μL of milliQ water and then heated to 100 °C for 3 min in a dry bath. The samples were centrifuged at 10 000 g for 5 min and the supernatant was used as a template to amplify a fragment that includes the V6, V7 and V8 variable regions of 16S rRNA gene. One μL of the template, 25 pmol of F_primer, 5’ GCACAAGCGGTGGAGCATGTGG 3’, and of the R_primer, 5’ GCCCGGGAACGTATTCACCG 3’, were mixed with 1x Taq buffer, 1.5 mmol of MgCl_2_, 0.2 mmol of each dNTP, and 1 U Taq DNA polymerase (ThermoFisher Scientific) in a final reaction of 25 μL. The amplification program was an initial denaturation at 94 °C for 5 min followed by 30 cycles at 94 °C for 45 sec, 50 °C for 45 sec, and 72 °C for 1 min, with a final extension of 10 min at 72 °C. Five μL of the final reaction was used as a template for the sequencing reaction. Sequencing reaction was done using Big Dye terminator v3.1 cycle sequencing kit (ThermoFisher Scientific) consisting of a 1x sequencing buffer, 25 pmol F_primer or R_primer and 2 μL of Big Dye in a final volume of 20 μL. The program used was an initial denaturation at 94 °C for 5 min, followed by 40 cycles at 94 °C for 30 sec, 50 °C for 30 sec, and 60 °C for 4 min. After the sequencing reaction, 80 μL of 70 % isopropanol was added and the reaction tube was centrifuged at 4000 g at 4 °C for 40 min. Then the supernatant was discarded, and the sample was resuspended in 20 μL of milliQ water and injected in an ABI PRISM 3130XL genetic analyzer (ThermoFisher Scientific). The obtained sequences were used to perform BLASTn (Altschul et al., 1990) searches against the Genbank/NCBI database (Benson et al., 2013) to identify most similar sequences.

### Genome sequencing, assembling and annotation

Bacterial cultures were grown in 2xTY broth (tryptone 16 g/L, yeast extract 10 g/L, and NaCl 5 g/L) at 37 °C for 18 h at 200 rpm. Genomic DNA purification was done using the Wizard Genomic DNA Purification Kit (Promega). DNA integrity was evaluated by 1 % agarose gel electrophoresis stained with SYBRSafe (Invitrogen) and by Bioanalyzer 2100 using Chips Agilent DNA 12000. DNA concentration and purity were estimated using a NanoDrop One/OneC Microvolume UV-Vis Spectrophotometer (ThermoFisher Scientific). Shotgun genomic library was prepared using the Nextera DNA Library Prep (Ilumina) with total DNA input of 20-35 ng. The resulting indexed DNA library was cleaned up with Agencourt AMPure XP beads (Beckman Coulter) and fragment size within the range of 200-700 bp were verified by running in the 2100 Bioanalyzer using Agilent High Sensitivity DNA chip. Fragment library quantification was performed with KAPA Library Quantification Kit. Genomic libraries prepared for each strain were pooled and subjected to a run using a an Ilumina MiSeq Reagent Kit v2 (2 × 250 cycles) which generated ~38 million raw paired-end reads with >75% of bases with quality score > 30.

The genome of strain PER-URP-08 was assembled with Discovar (v. 52488) (Weisenfeld et al., 2014). The genomes of strains PER-URP-12 and PER-URP-17 were assembled with A5 (v. 20160825) (Coil et al., 2015). Both software have adapters trimming and read quality checking as part of their respective assembly processes. The tool Medusa (Bosi et al., 2015) was used to generate final genome scaffolds using three sets of five reference genomes, one for each of the genome assemblies (Table S1). The final genome assemblies were submitted to the IMG/M (Chen et al., 2021) and to the NCBI (Benson et al., 2013; Tatusova et al., 2016) for automatic annotation.

### Phylogenetic analyses and identification of nitrilases

Annotated genomes belonging to *Bacillus pumilus, Bacillus safensis* or *Bacillus altitudinis* species in the category of “Chromosome”, “Scaffold” or “Complete” were downloaded from the Genbank/NCBI (Benson et al., 2013). Using the software cd-hit (Fu et al., 2012; Li & Godzik, 2006) we identified coding sequences that are not duplicated and present in all the genomes (core genes). A total of 1766 core genes with more than 80 % identity and at least 90 % coverage were used in the analysis. Core genes were aligned using MAFFT with the FFT-NS-2 algorithm (Katoh & Standley, 2013). The resulting alignments were concatenated and used to calculate a distance matrix based on identity using Biopython (Cock et al., 2009). Phylogenetic inference by maximum likelihood was done using the concatenated alignments as the input and IQ-TREE2 (Minh et al., 2020) with the evolution model GTR+F+R3, ultrafast bootstrap 1000 (Hoang et al., 2018), and 1000 initial trees.

IMG/M tools (Chen et al., 2021) were used to identify nitrilases genes in the annotated genomes. Genes encoding the CN_hydrolase domain (PFAM code PF00795) were selected and checked regarding the genomic context and the related literature.

### Analysis of CynD sequences from *Bacillus pumilus* group genomes

Protein sequence annotations from genomes belonging to *Bacillus pumilus, Bacillus safensis* or *Bacillus altitudinis* in the category of “Chromosome”, “Scaffold” or “Complete” were downloaded from GenBank/NCBI (Benson et al., 2013) and used to construct a local database. We ran a BLASTp search (Altschul et al., 1990) using the query sequence AAN77004.1 against the constructed local database, and sequences with more than 90 % identity and 100 % coverage were identified as CynD orthologs. These sequences were aligned using MAFFT with the L-INS-I algorithm (Katoh & Standley, 2013). The resulting alignment was used as an input for the phylogenetic inference by maximum likelihood using IQ-TREE2 (Minh et al., 2020) with the evolution model JTTDCMut+I (Kosiol & Goldman, 2005), ultrafast bootstrap 1000 (Hoang et al., 2018), 1000 initial trees and -allnni option.

### Cloning, expression and purification of CynD

The coding sequence for CynD was amplified from genomic DNA of strain PER-URP-08 using the primers (restriction sites appear in uppercase): F_CynD (5’ tttCATATGatgacaagtatttacccgaagtttc 3’), and R_CynD (5’ tttCTCGAGcactttttcttcaagcaaccc 3’) and cloned in the NdeI and XhoI sites of pET-28 plasmid. Then, this plasmid was used as a template to amplify the CynD coding sequence with a c-terminal 6x-His tag using the primers F_CynD and R_2_CynD (5’ tttGAATTCagtggtggtggtggtggtg 3’) and cloned in the NdeI and EcoRI sites of pET-11 plasmid.

To express CynD protein, we used the *Escherichia coli* BL21(DE3) pLysS strain, induced by 0.3 mM of Isopropyl β-D-1-thiogalactopyranoside for 23 h at 18°C. The cells were lysed by sonication using a lysis buffer (100 mM Tris-HCl pH 8.0, 100 mM NaCl, 50 mM Imidazole) and the suspension was clarified by centrifugation (13000 g). The supernatant was loaded onto Ni-NTA affinity resin (His-trap chelating 5 mL column), washed with 10 volumes of lysis buffer, and eluted with a gradient of 50 - 500 mM Imidazole in 20 mM Tris-HCl pH 8.0, 100 mM NaCl. The eluted fractions were further purified by size exclusion chromatography using a Superdex pg 200 16/600 column and 20 mM Tris-HCl pH 8.0, 100 mM NaCl as running buffer. The eluted fractions were examined for purity by SDS-PAGE and fractions containing pure protein were concentrated in Amicon Ultra-15 Centrifugal filter units.

### Enzymatic assays of recombinant CynD

For the determination of Km and Vmax, enzymatic activity of recombinant CynD was measured at pH 8.0 using the Ammonia Assay Kit (Sigma-Aldrich). A concentration of 500 nM of CynD was used in all reactions with the following cyanide concentrations: 0.39, 0.625, 0.78, 1.25, 1.56, 2.5, 3.125, 5, 6.25, 12.5, 25 mM, with a final volume of 111 uL at 30 °C. Measurements were taken on a plate reader, at 340 nm every 20 sec. Calculations of ammonia concentrations were carried out according to the manufacturer description.

To test in which optimal pH for CynD activity, reagent solutions were prepared (40 mM NaCN, 100 mM NaCl and 200 mM Tris-HCl or N-cyclohexyl-3-aminopropanesulfonic acid (CAPS) at pH 8, 9 or 10, 11, respectively). Then, we added 5 μL of CynD in 100 mM NaCl, Tris-HCl pH 8 to 45 μL of the reagent solutions to obtain a final concentration of CynD of 0, 5, 10, 15, or 20 μM. The reactions were incubated for 10 min at 37 °C. After that, 100 μL of picric acid 5 mg/mL, 0.25 M Na_2_CO_3_ was added, and the reactions were incubated at 99 °C for 6 min. Next, 30 μL of this reaction was transferred to a 96-well plate and absorbance at 520 nm was recorded. Final cyanide concentration was estimated based on calibration curves with cyanide concentrations between 0 – 40 mM.

### Size Exclusion Chromatography coupled to Multi-Angle Light Scattering (SEC-MALS)

SEC-MALS analysis was used to determine the oligomeric state of recombinant CynD. Molar mass analysis was done in 100 mM NaCl and 20 mM Tris-HCl or CAPS at pH 8, 9 or 10, 11, respectively. Protein samples (100 μL injection of 3.47 mg/mL (89.36 μM) CynD) were separated using a Superdex 200 increase 10/300 GL coupled to a MiniDAWN TREOS multi-angle light scattering system and an Optilab rEX refractive index detector. Data analysis was performed using the Astra Software package, version 7.1.1 (Wyatt Technology Corp.).

### Transmission electron microscopy (TEM)

Ultra-thin carbon layer on lacey carbon-coated copper grids were negatively charged by a glow discharge of 25 sec at 15 mA. Four microliters of purified recombinant CynD in 20 mM Tris-HCl pH 8.0 and 100 mM NaCl in different concentrations (3.25 mg/mL or 1.625 mg/mL) were placed in the negative charged carbon-coated copper grid for 1 minute. The grids were washed twice with MilliQ water and then stained with 2 % uranyl acetate for 30 secs before air drying. Electron micrographs images were obtained using a JEOL JEM 2100 transmission electron microscope equipped with a Gatan ORIUS CCD detector at the Institute of Chemistry of the University of Sao Paulo.

### RT-qPCR to evaluate *in vivo* induction of *cynD* by cyanide

*Bacillus* strains were grown in meat broth (meat extract 1 g/L, yeast extract 2 g/L, peptone 5 g/L, NaCl 5g/L, MnCl_2_ 10 mg/L) during 12 h at 30 °C, 200 rpm. One mL of the culture was centrifuged at 500 xg for 1 min, the supernatant was transferred to a clean tube and this tube was centrifuged at 8 000 xg for 3 min. The pellet was resuspended in 1 mL of NaCN ([CN^−^] 100 ppm) in milliQ water. Controls were resuspended in 1 mL milliQ water without NaCN. The tubes were incubated without agitation at 30 °C for 4 h and 100 μL were retrieved to measure cyanide concentration by the picric acid method (Williams & Edwards, 1980). Nine hundred μL was centrifuged, and the bacterial pellet was used immediately for total RNA extraction.

Total RNA extraction was done using Trizol-chloroform protocol. Briefly, bacterial pellets were treated with 100 μL of lysozyme 3 mg/mL at 37 °C for 30 min, and extraction was done using a mixture of 5:1 trizol:chloroform. After the extraction, the phase containing RNA was separated and the RNA was precipitated using isopropanol. RNA pellet was washed twice with 75 % ethanol and finally resuspended in 20 μL of Tris 20 mM-DEPC. Total RNA concentration and purity were estimated in a NanoDrop™ One/OneC Microvolume UV-Vis Spectrophotometer (ThermoFisher Scientific) and the integrity was evaluated in a 2100 Bioanalyzer using an Agilent RNA 6000 Pico chip. After DNase treatment, the samples were subjected to PCR to verify the absence of DNA contamination. cDNA synthesis was performed with 1 μg of the RNA and Thermo Scientific H Minus First Strand cDNA Synthesis kit. cDNA synthesis was verified by PCR and electrophoresis.

Amplification efficiency of the primers used in the RT-qPCR were verified using 300 nM of each primer and a 2-fold dilution series of the cDNA to generate a standard curve composed of 4 concentrations as follows: 62.5, 31.25, 15.625, and 7.8125 ng/μL. Each dilution reaction was performed in triplicate using the Maxima SYBR Green/ROX qPCR Master Mix kit (ThermoFisher Scientific) following the manufacturer instructions in a QuantStudio 3 equipment (ThermoFisher Scientific). Primers for the normalizing gene *rpsJ* (F_rpsJ 5’ TGAAACGGCTAAGCGTTCTG 3’, R_rpsJ 5’ ACGCATCTCGAATTGCTCAC 3’), and for the nitrilases *cynD* (F_cynD 5’ TGCCCAAAATGAGCAGGTAC 3’, R_cynD 5’ AAATGTCTGTGTCGCGATGG 3’) and *ykrU* (F_ykrU 5’ TTGGTGCGATGATTTGCTAT 3’, R_ykrU 5’ GTGTCTCTGCTTGTGCCTGT 3’) were tested for efficiency. The amplification efficiency of the qPCR reaction was calculated through the slope of the cDNA curve obtained for each primer pair.

Since primer pairs have showed similar efficiency, (*ykrU* = 119.108 %, *cynD* = 108.385 %, *rpsJ* = 104.55 %), we performed each qPCR assay in technical triplicates using 15.625 ng/μL of cDNA and the kit Maxima SYBR Green/ROX qPCR Master Mix (ThermoFisher Scientific) in a QuantStudio 3 equipment (ThermoFisher Scientific). ΔΔCT values were calculated in absence or presence of cyanide for the nitrilase genes *ykrU* and *cynD* using *rpsJ* as the normalizing gene. Three biological replicates were performed.

## RESULTS AND DISCUSSION

### Three *Bacillus spp*. isolates with capacity of cyanide degradation

Several colonies were obtained after selective enrichment in cyanide containing media of water in contact with mine tailing from a river near Casapalca and La Oroya mines located in San Mateo de Huanchor, Lima - Peru. Twenty colonies were screened for the ability to degrade cyanide (Table S2) and three colonies with the greatest efficiency in cyanide degradation (isolates 8, 12, and 17) were selected for further studies (Table S2).

Sequencing of the V6, V7, and V8 variable regions of 16S rRNA gene of the three selected isolates and analysis by BLAST showed that they belong to the genus *Bacillus* (Table S3). Isolates 12 and 17 were identified as *Bacillus licheniformis* and *Bacillus subtilis*, respectively (Table S3) and were named *Bacillus licheniformis* PER-URP-12 and *Bacillus subtilis* PER-URP-17. Isolate 8 was classified as a member of the *Bacillus pumilus* group based on the 16S rRNA gene sequence (Table S3). However it was not possible to discriminate among the different species in the *Bacillus pumilus* group (Liu et al., 2013) and as such this isolated was provisionally named *Bacillus sp*. PER-URP-08.

The three strains were then sequenced in order to obtain a more accurate taxonomical classification as well as to gain insights about possible routes of cyanide degradation in the three strains under study. Table S4 shows a summary of assembly and annotation metrics of these genomes.

### *Bacillus sp*. PER-URP-08 is classified as *Bacillus safensis* based on core-genome comparisons

We performed a genome-wide comparative analysis of *Bacillus sp*. PER-URP-08 with 132 genomes of species from the *Bacillus pumilus* group retrieved from the GenBank/NCBI database (Benson et al., 2013) and identified 1766 coding sequences present in all the genomes (core genes). An identity matrix based on an alignment of these core genes showed three well defined branches and two genomes that do not belong to any of these three branches (Fig. 1A).

**Figure 1.**
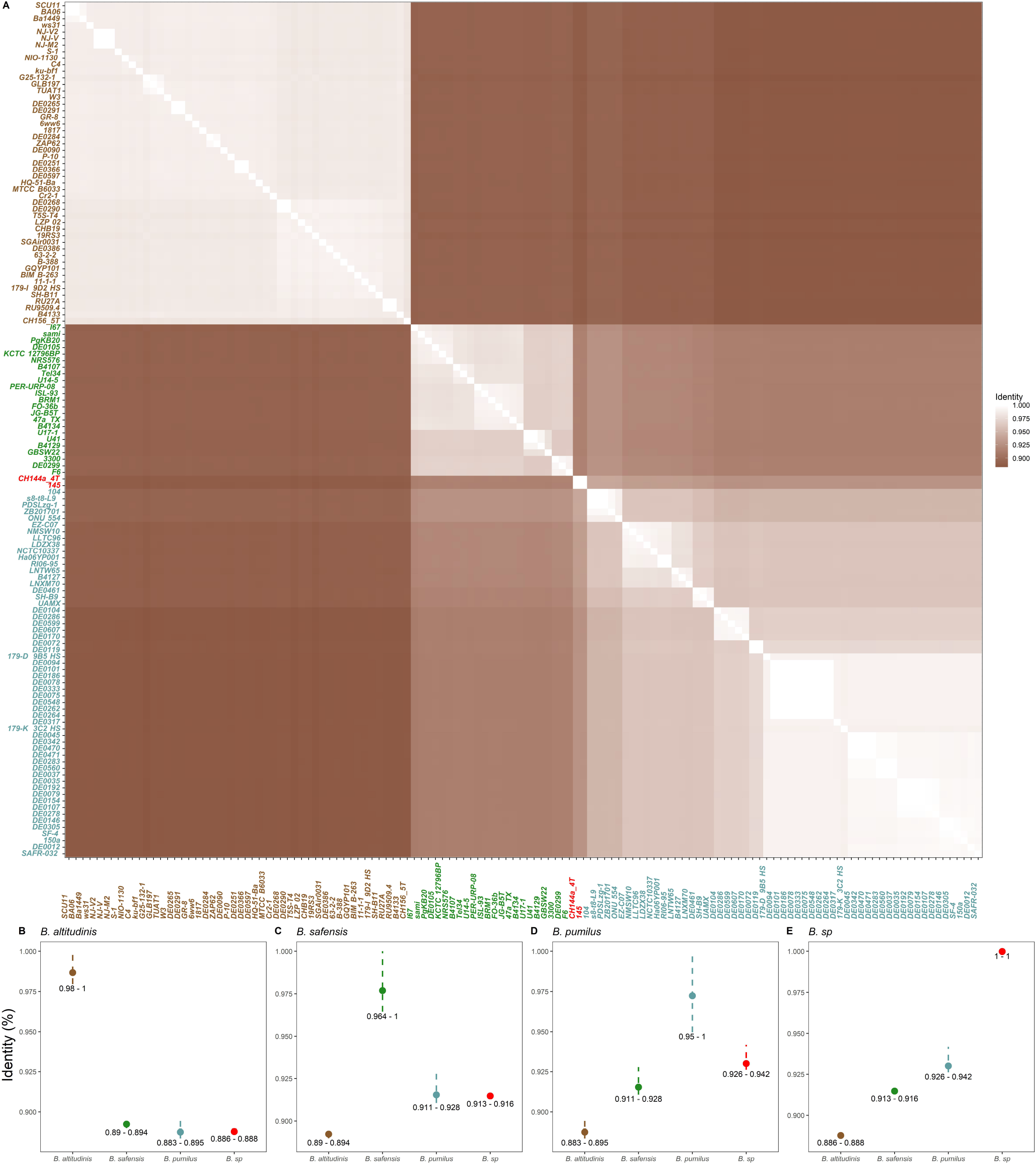
Core genome identity matrix to classified genomes of *Bacillus pumilus* group genomes. A) An identity matrix of 132 core genomes of *Bacillus pumilus* group showing delimitations between three species: *Bacillus altitudinis* (brown names), *Bacillus safensis* (green names), *Bacillus pumilus* (blue names). Two core genomes (red names) appear outside of these three species. B – E) Plots showing the range of identity when compare *B. altitudinis* (B), *B. safensis* (C), *B. pumilus* (D) or *B. sp* (E) with itself or with other groups.

Branch 1 (Fig. 1A, brown names) contains several strains already characterized as *Bacillus altitudinis* by different methods (for instance: BA06, ku-bf1, B-388 (X. Fu et al., 2021)) and also 4 strains (TUAT1, MTCB 6033, SH-B11 and C4) previously annotated as *Bacillus pumilus*. However, our analysis clearly demonstrates that they belong to *Bacillus altitudinis* and therefore require reclassification (Table S5) as previously suggested (Espariz et al., 2016; X. Fu et al., 2021). The core genes within the *Bacillus altitudinis* branch share more than 0.98 identity whereas they share less than 0.895 identity with core genomes of the other two branches (Fig. 1B).

Identity of core genes in branch 2 is greater than 0.96, and this branch is more related to branch 3 (*Bacillus pumilus, see below*) than to branch 1 (*Bacillus altitudinis*) (Fig. 1C). Branch 2 (Fig. 1A, green names) contains the *Bacillus safensis* type strain FO-36b (Satomi et al., 2006) as well as other strains already classified as *Bacillus safensis* such as B4107, B4134, and B4129 (Espariz et al., 2016). *Bacillus sp*. PER-URP-08 appeared inside this branch very near to the type strain FO-36b (99.2 % identity) (Fig. 1A) and so will be named *Bacillus safensis* PER-URP-08 from here on.

Branch 3 (Fig. 1A, blue names) contains the SAFR-032 strain that was the first completely sequenced genome of *Bacillus pumilus* (Gioia et al., 2007; Stepanov et al., 2016). This branch 3 appears to be more heterogeneous than the other two branches (*Bacillus altitudinis* and *Bacillus safensis*) with more than 0.95 identity of the core genes of this branch (Fig. 1D). Additionally, two genomes isolated from Mexico (CH144a_4T and 145) share less than 0.95 identity with the branch 3 and even less with branches 1 and 2 (Fig. 1E). The fact that these two genomes share less than 0.95 identity with all the other genomes in the analysis (Fig. 1E) indicates that CH144a_4T and 145 strains should be classified as different species outside the *Bacillus pumilus* group.

### A cyanide dihydratase is likely the responsible for cyanide degradation in *B. safensis* PER-URP-08

To gain insight regarding the enzymes responsible for cyanide metabolism in the strains *B. safensis* PER-URP-08, *B. licheniformis* PER-URP-12, and *B. subtilis* PER-URP-17, we first searched for genes coding for proteins related to nitrilases. The PFAM database annotates homologs of nitrilases as CN_hydrolases under the PFAM code PF00795. Using IMG/M system tools (Chen et al., 2021), we determined the presence of three, two, and two proteins containing CN_hydrolase domains in *B. safensis* PER-URP-08, *B. licheniformis* PER-URP-12, and *B. subtilis* PER-URP-17, respectively (Fig. S1). Both *B. licheniformis* PER-URP-12 and *B. subtilis* PER-URP-17 present the genes *yhcX* (NCBI locus tags: EGI08_RS06285 and EGI09_16505, respectively) and *mtnU* (EGI08_RS08970 and EGI09_01680, respectively). *YhcX* is probably involved in the degradation of indole-3-acetonitrile, a sub product of tryptophan metabolism (Idris et al., 2007) (Fig. S1). On the other hand, MtnU has been described as a possible enzyme catalyzing the conversion of alpha-ketoglutaramate to alpha-ketoglutarate involved in the metabolism of methionine (Ellens et al., 2015; Sekowska & Danchin, 2002) (Fig. S1). None of the enzymes with a CN_hydrolase domain in *B. licheniformis* PER-URP-12 and *B. subtilis* PER-URP-17 appears to be responsible for cyanide degradation. However, apart from these proteins, *Bacillus* and other genera present proteins with rhodanese domains (PFAM codes PF12368 and PF00581) (Table S6) that are able to convert cyanide to thiocyanate (Cipollone et al., 2006; Itakorode et al., 2019). Thus, it is likely that those rhodanese enzymes are responsible for the degradation of cyanide by *B. licheniformis* PER-URP-12 and *B. subtilis* PER-URP-17. Further studies are necessary to test this hypothesis.

*B. safensis* PER-URP-08 presents *yhcX* (EGI07_01665) but not *mtnU*. In addition, this strain carries two other proteins containing a CN_hydrolase domain, EGI07_17510 and CynD (EGI07_08135). EGI07_17510 is a protein of unknown function whereas CynD homologs (Fig. S2) hydrolyzes cyanide to produce ammonia and formic acid (Dash et al., 2009; Ibrahim et al., 2015). We therefore carried out a series of experiments to test the hypothesis that CynD is the enzyme responsible for cyanide degradation in *B. safensis* PER-URP-08.

### C-terminal residues differentiate CynD from *B. pumilus* and *B. safensis*

We first constructed a maximum likelihood (ML) phylogenetic tree based on the 132 core genomes of strains from *Bacillus pumilus* group (Fig. 2A) and searched for orthologs of CynD in the strains present in the ML tree (see Methods for details of the search). The ML tree confirmed the three branches identified above (Fig. 1A) and that two genomes (CH144a_4T and 145) do not belong to any of these branches (Fig. 2A). Intriguingly, CynD-encoding sequences were found in some representatives of *B. pumilus* (44 out of 56) and *B. safensis* (19 out of 23) but not in *B. altitudinis*. Three monophyletic *B. pumilus* and one monophyletic *B. safensis* clades lack CynD (Fig. 2A). This could be due to processes of gene gain and/or loss in the strains, and further studies are necessary to distinguish between these or other possibilities. It is also possible that some *cynD* genes were no sequenced in some incomplete genomes.

**Figure 2.**
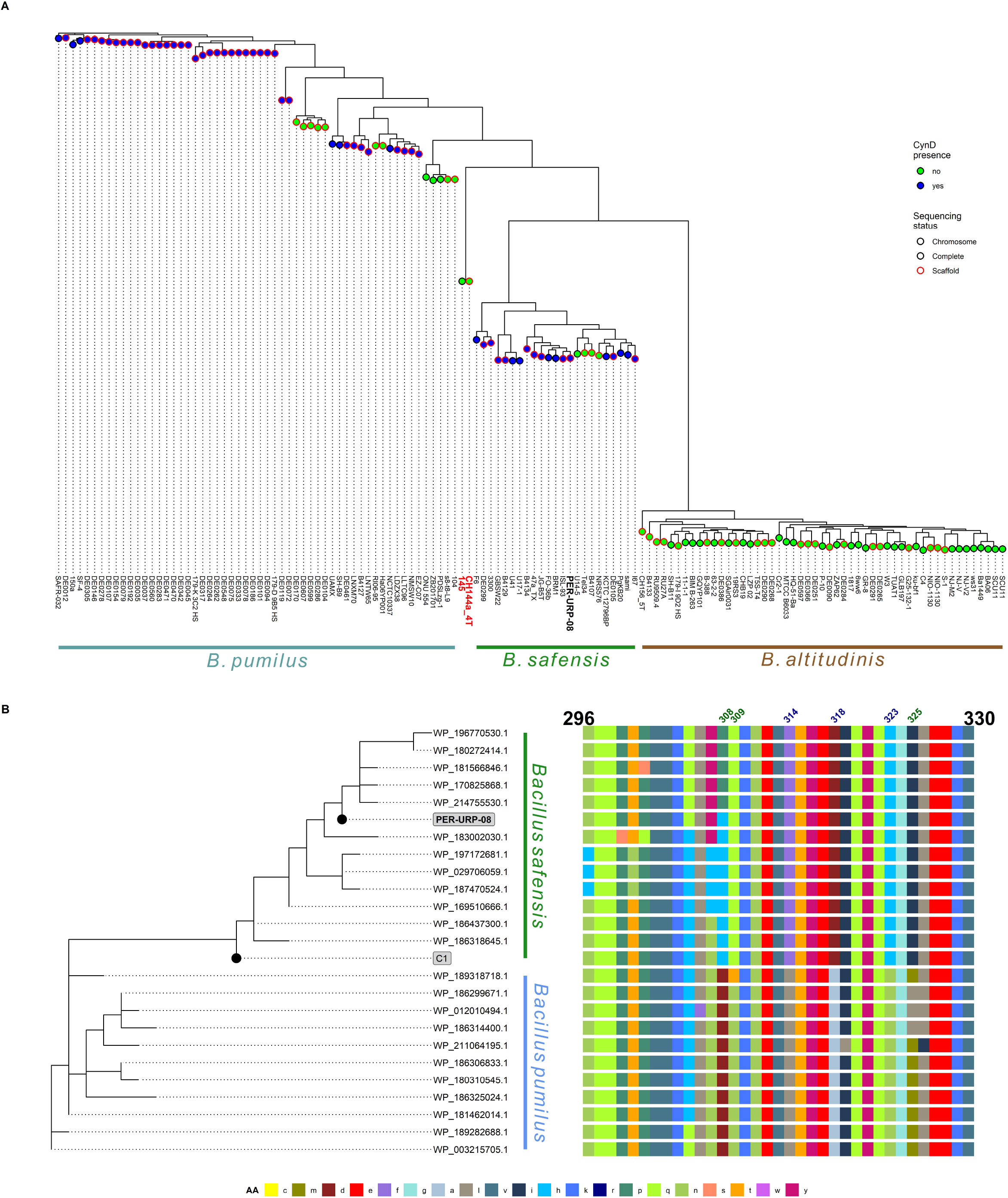
CynD is present in some genomes of *B. pumilus* and *B. safensis* and they are mainly differentiated by C-terminal residues. A) Maximum likelihood tree of core genomes of 132 *Bacillus pumilus* group strains showing separation between three species. Color of the circles represent absence (green) or presence (blue) of CynD homologue in the genome. Circles with black and red borders represent complete genomes (“chromosome” or “complete” sequencing status in NCBI) and possibly not complete genomes (“scaffold” sequencing status in NCBI). B) Maximum likelihood tree of full-length CynD sequences associated to and alignment of their C-terminal region (residues 296 to 330). Showed in number blue or green are the positions that are completely conserved in *Bacillus safensis* or *B. pumilus*, respectively.

Next, we identified twenty-three different sequences of CynD in the 132 genomes (Table S7) and a ML phylogenetic tree based on aminoacid sequences was constructed, including the sequences of the CynD from strain C1 (CynD_C1_) (accession id: AAN77004.1) and of the CynD from *B. safensis* PER-URP-08 (CynD_PER-URP-08_). A clear separation between CynD from *B. safensis* and from *B. pumilus* could be observed in the ML tree (Fig. 2B). Interestingly, CynD_C1_ appear more related to the *B. safensis* group (Fig. 2B). Due to the several taxonomic misclassifications of strains belonging to the *Bacillus pumilus* group (as reported here and by others (Espariz et al., 2016; X. Fu et al., 2021; Liu et al., 2013)), it is likely that strain C1 truly belongs to a *B. safensis* species; however, complete genome of C1 is not available to confirm this hypothesis.

The most variable region in the nitrilase protein family is the C-terminal (Benedik & Sewell, 2018; Thuku et al., 2009). Thus, we associated a phylogenetic tree obtained from the full-length sequences of identified CynDs homologs to an alignment of the C-terminal region (residues 296 to 330) (Fig. 2B). Residues F314, D318, H323 in *B. safensis* CynD are L314, A318, and N323 in the *B. pumilus* protein. Other residues can vary in one of the species but are strictly conserved in the other, for instance, residues Q309 and I325 in *B. safensis* are T309 or N309 and M325 or L325 in *B. pumilus*. Residue 308 can be P or M in *B. safensis* but is strictly D in *B. pumilus* (Fig. 2B). CynD_C1_ has the aminoacids strictly conserved in *B. safensis* supporting the conclusion that C1 belongs to *B. safensis* species. Furthermore, residue 27, outside the C-terminal, is E in *B. safensis* and strain C1 but Q in *B. pumilus*.

### CynD from *B. safensis* PER-URP-08 it is still active at pH 9

We then went on to characterize some biochemical properties of CynD_PER-URP-08_. First, we cloned and expressed recombinant CynD_PER-URP-08_ in *E. coli* and determined the basic kinetic constants of the purified recombinant enzyme. Although CynDs are known to be able to adopt different oligomeric states, no evidence of cooperativity was observed in our enzymatic assays (Fig. 3A). Instead, a simple Michaelis-Menten model fit the experimental data adequately. K_m_ and k_cat_ estimated using this model were 1.93 mM and 6.85 s^−1^ (Fig. 3A, S3). The K_m_ value is similar to that previously reported for CynD_C1_ and from other species (2.56 mM to 7.3 mM) similar but k_cat_ is lower (433 s^−1^ to 61600 s^−1^) (Crum et al., 2015; Crum et al., 2016; Jandhyala et al., 2005; Vargas-Serna et al., 2020).

**Figure 3.**
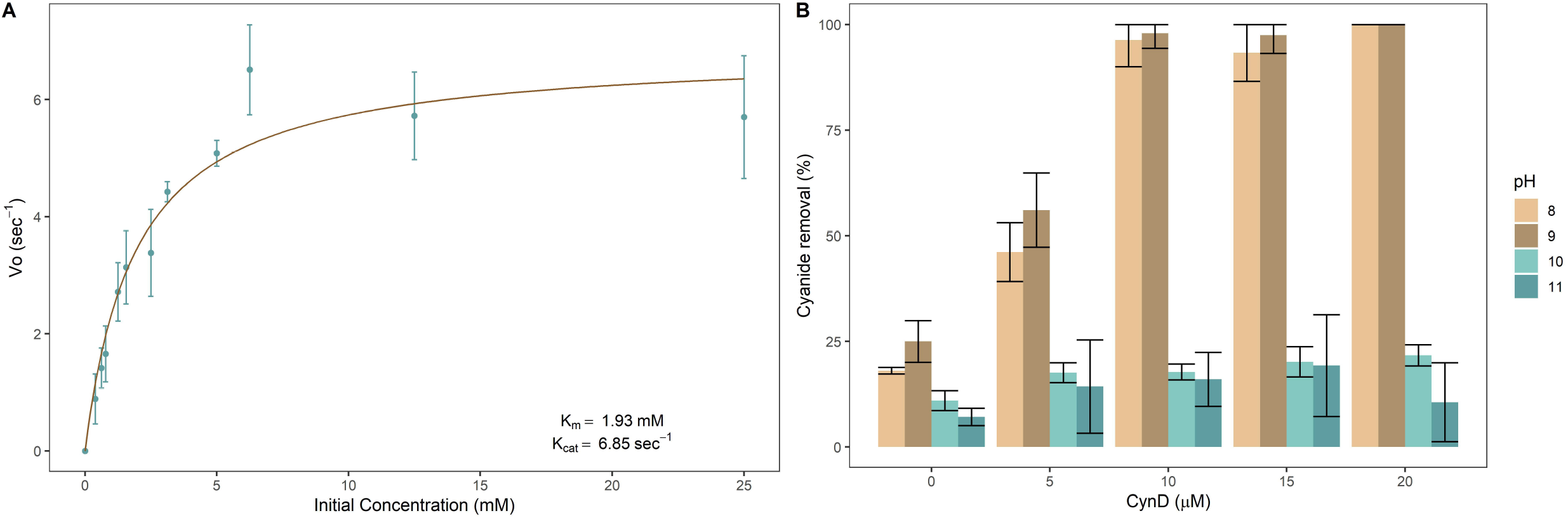
CynD_PER-URP-08_ have similar kinetic constants to other CynD homologues and is still active up to pH 9. A) Plot of CynD_PER-URP-08_ Initial velocity (Vo) versus initial concentration of cyanide adjusted to the Michaelis Menten equation. Km and Kcat constants calculated assuming this model are shown in the graphic. B) Percentage of cyanide removal in different pHs using different CynD_PER-URP-08_ concentrations. CynD_PER-URP-08_ showed considerable activity in pH 8 and 9 but not in 10 and 11.

Due to the volatility of hydrogen cyanide in its protonated HCN state and its pKa of 9.2 (Brüger et al., 2018), bioremediation processes should preferably be carried out at or above pH 9. To test if CynD_PER-URP-08_ is active at pHs greater than 8, we tested its activity at pH 9, 10, and 11. Figure 3B shows that recombinant CynD_PER-URP-08_ carrying a C-terminal 6x-His tag is active up to pH 9 and inactive at pH 10 and 11. Other wild-type CynDs have been shown to be active only up to pH 8 (Crum et al., 2016; Jandhyala et al., 2005) and CynD_C1_ with C-terminal 6x-His tag had its activity compromised at pH 9 (Vargas-Serna et al., 2020). The CynD_C1_ and CynD_PER-URP-08_ sequences only differ at five positions: are I18V, S25T, E155D, H305Q and N307Y (first letter correspond to CynD_C1_) with the last two substitutions H305Q and N307Y near the C-terminus.

Other studies were able to generate active versions of CynD active at pH 9 by introducing mutations in some conserved positions (K93R; Q86R, E96G, D254G) or by replacing the C-terminal from CynD_C1_ with the C-terminal from CynD from *Pseudomonas* stutzeri (Crum et al., 2015; Wang et al., 2012) (note that wild-type CynD from *P. stutzeri* has not been tested at pH 9).

### Alkaline pH reduces the degree of oligomerization of CynD_PER-URP-08_

The oligomerization state of nitrilases have been associated with enzyme activity and stability (Crum et al., 2015; Crum et al., 2015; Crum et al., 2016; Martínková et al., 2015; Park et al., 2016; Wang et al., 2012). In the case of CynDs of CynD_C1_ and CynD from *P. stutzeri*, mutations in the C-terminal region decrease oligomerization (M. Crum et al., 2016; M. A. N. Crum et al., 2015; Wang et al., 2012). The C-terminal of nitrilases stabilizes the spiral structure through crisscrossed beta sheets in the center of the oligomer (Mulelu et al., 2019; Thuku et al., 2009). Also, pH has been shown to promote higher order oligomerization states of CynDs (D. Jandhyala et al., 2003; Wang et al., 2012); however, the effects of pH greater than 9 have not been reported.

Since, CynD_PER-URP-08_ has differences in C-terminal with respect to other CynDs we used SEC-MALS to compare the oligomerization states of CynD_PER-URP-08_ at different pHs. As expected, pHs higher than 8 results in smaller sized oligomers. At pH 11 the monomer (38.5 kDa) is the predominant species (Fig. 4A), whereas pH 10 and 9 presented oligomeric states ranging from ~3-mer to ~5-mer (pH 10, 100.85 to 176.34 kDa) and ~4-mer to ~6-mer (pH 9, 133.19 to 226.99 kDa) (Fig. 4B-C). Furthermore, CynD_PER-URP-08_ presented oligomers ranging from ~24-mer to ~48-mer (918.31 to 1851.39 kDa) at pH 8 (Fig. 4D) in contrast to what was reported for CynD_C1_ at pH 8 which forms an 18-mer spiral (D. Jandhyala et al., 2003). These differences could be a result of the differences in aminoacid sequence between CynD_PER-URP-08_ and CynD_C1_ or due to the presence of the C-terminal 6x-His tag in CynD PER-URP-08. Experiments with CynD_C1_ were carried out with untagged protein or with protein carrying an N-terminal 6x-His tag (Crum et al., 2015; Jandhyala et al., 2003; Park et al., 2016; Wang et al., 2012). Electron micrographs of negatively stained CynD_PER-URP-08_ at pH 8 showed spirals of different sizes supporting the conclusion that CynD_PER-URP-08_ at this pH is adopts a range of different oligomerization states (Fig. 4E).

**Figure 4.**
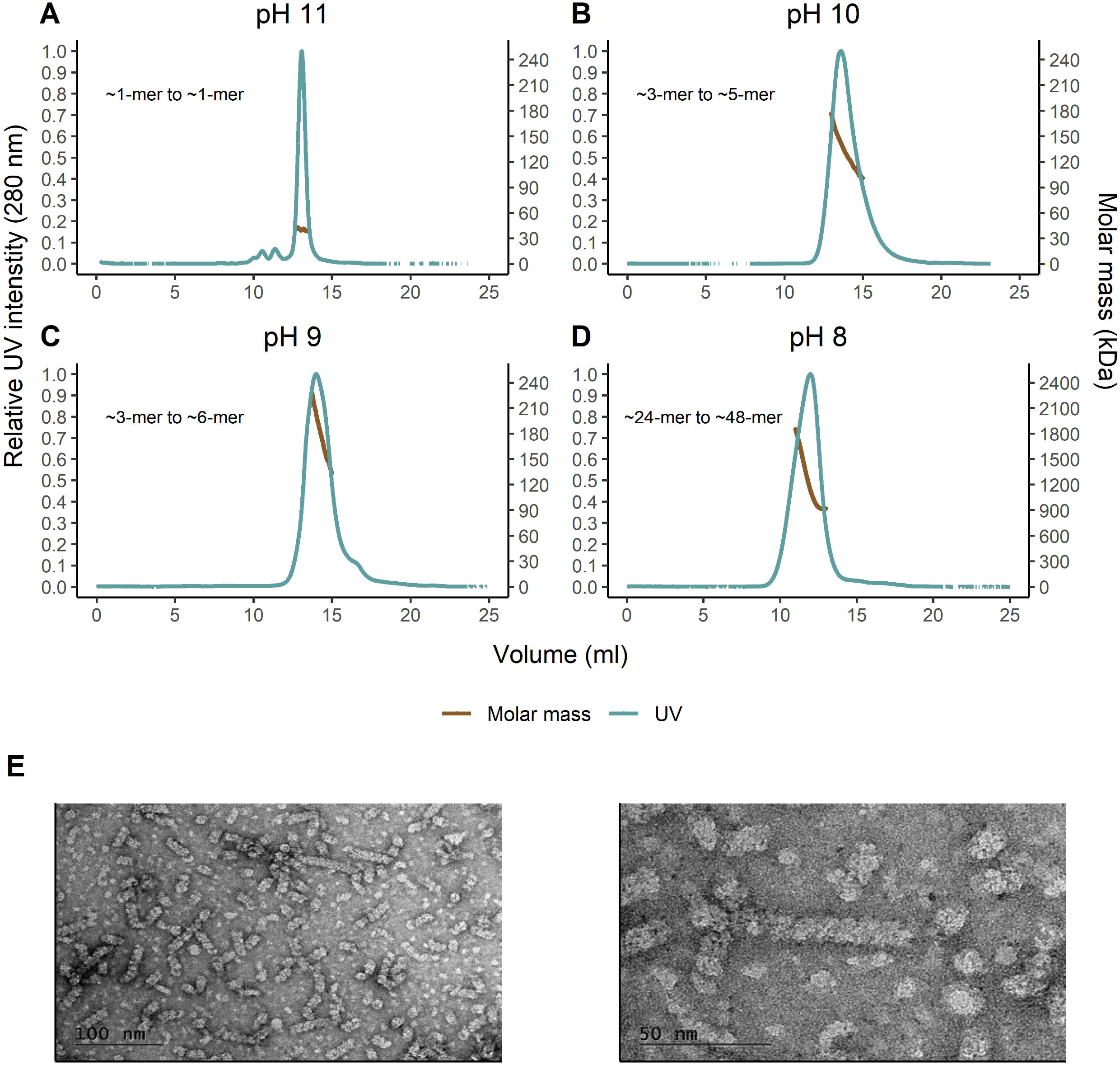
SEC-MALS of CynD_PER-URP-08_ showed that higher pHs reduced its oligomerization states and TEM showed that CynD_PER-URP-08_ presents a helical structure. A-D) Plot of UV intensity/molar mass for CynD_PER-URP-08_ in different pHs. A pattern of decrease the oligomeric state while increasing the pH was observed. E) TEM micrographs at pH 8 in two different magnifications (right and left) showed helical structures of CynD_PER-URP-08_.

### Expression of CynD_PER-URP-08_ from *B. safensis* PER-URP-08 is induced in the presence of cyanide

Some previous studies have considered the possibility that CynD gene expression is regulated by cyanide, but this point remains unclear (D. Jandhyala et al., 2003). To address this question, we exposed *B. safensis* PER-URP-08 to 100 ppm CN^−^ (in the form of 38.5 mM NaCN) at 30 °C for 4 h without agitation and the mRNA levels of *cynD* were measured and compared with the levels observed in cells grown in the absence of CN^−^.

We observed a 6.7-fold increase in expression of *cynD* in the presence of cyanide (Fig. 5). To evaluate if this overexpression is specific for *cynD* nitrilase and not to other nitrilases of *B. safensis* PER-URP-08, we also measured the mRNA levels of *ykrU* that also possesses a CN_hydrolase domain. We did not observe differences in *ykrU* expression in the presence and absence of cyanide. To our knowledge, this is the first report showing induction in the expression of *cynD* in the presence of cyanide. This could possibly be a physiological response of the bacteria in order to protect itself from the toxic effects of the compound, but further studies are necessary to more fully understand the molecular mechanisms behind this response.

**Figure 5.**
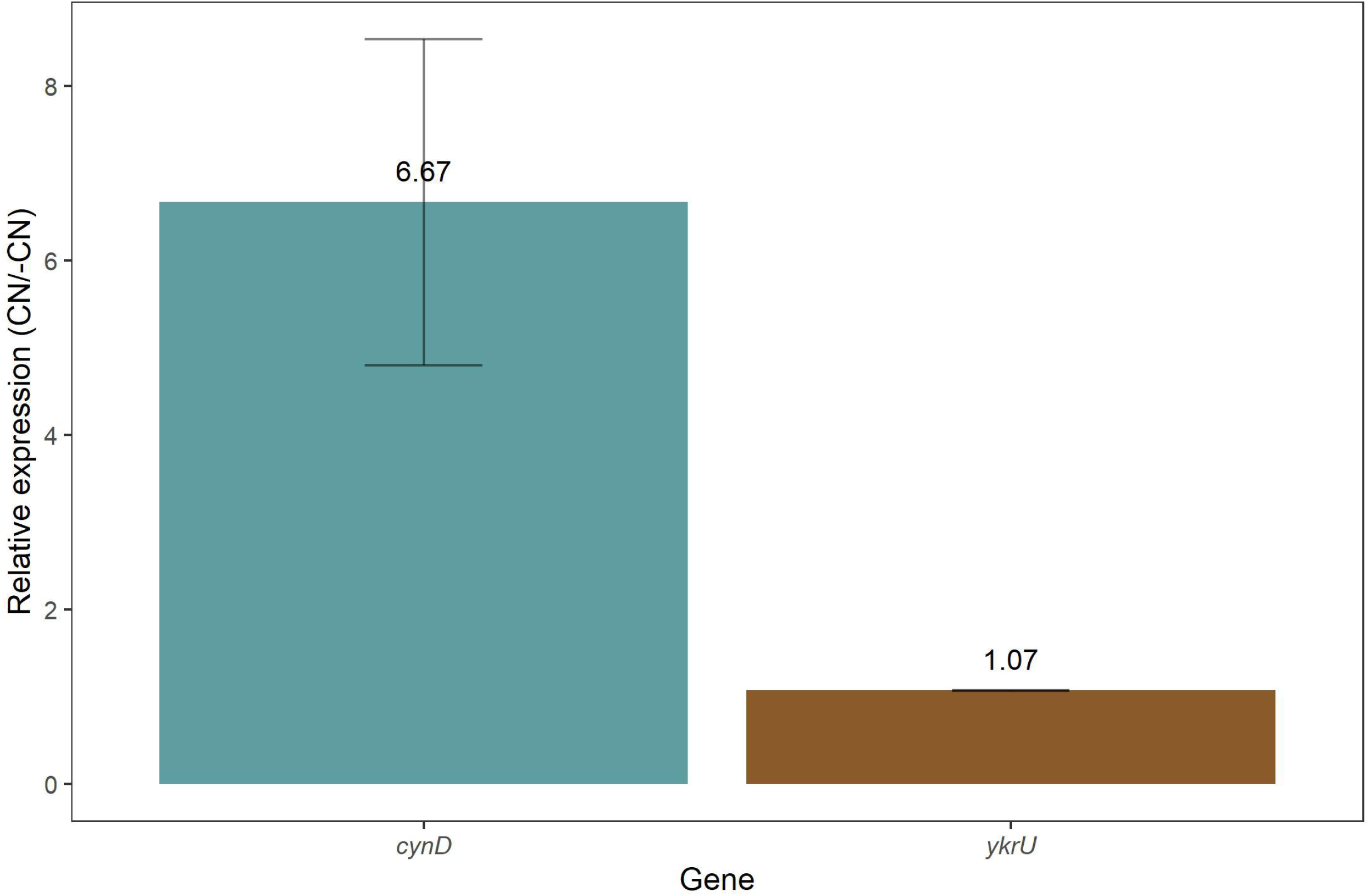
*cynD*_PER-URP-08_ but not *ykrU* is induced in the presence of cyanide. Relative expression measured by RT-qPCR showed that when *Bacillus safensis* is in presence of cyanide the RNA levels of *cynD* are 6.67-fold greater than when in absence of cyanide. In contrast, other nitrilase gene (*ykrU*) have the same RNA levels in presence or absence of cyanide.

## CONCLUSIONS

Here we report the isolation and the genome sequences of three cyanide-degrading *Bacillus* strains obtained from water in contact with mine tailings in Lima – Peru. They were phylogenetically classified and named *Bacillus licheniformis* PER-URP-12, *Bacillus subtilis* PER-URP-17 and *Bacillus safensis* PER-URP-08. Comparative genomic analyses indicate that some strains currently classified as *B. pumilus* with publicly available genomes should be reclassified as *Bacillus altitudinis* (strains TUAT1, MTCB 6033, SH-B11, and C4). Furthermore, we propose that strains CH144a_4T and 145 should be classified belonging a new species distinct from *B. pumilus, B.safensis*, or *B. altitudinis*.

We propose that in *B. licheniformis* PER-URP-12 and *B. subtilis* PER-URP-17 rhodaneses (table S6) are possibly the enzymes that confer cyanide degradation capabilities to these strains. In the case of *B. safensis* PER-URP-08, we suggest that EGI07_08135 codes for an ortholog of cyanide dihydratase CynD that imparts the cyanide-degradation ability to this strain.

We found that while no *B. altitudinis* strains code for CynD orthologs, some *B. pumilus* and *B. safensis* strains present CynD orthologous sequences. CynD from *B. pumilus* and *B. safensis* have high identity (> 97%), however conserved differences in the C-terminus allow us to differentiate between CynD from *B. safensis* or *B. pumilus* (at least in the analyzed genomes). Additionally, sequence analysis of the previously described CynD from strain C1 (CynD_C1_), named as *B. pumilus* CynD in the literature, is more closely related to CynDs from *B. safensis* than from *B. pumilus*. Thus, indicating that CynD_C1_ is a representative of *B. pumilus* CynDs.

We characterized some aspects of CynD from *B. safensis* PER-URP-08 (CynD_PER-URP-08_) corroborating what was described for CynDs from other species and adding new knowledge about these enzymes. First, enzymatic assays with CynD_PER-URP-08_ found no evidence of cooperativity despite the known oligomerization patterns of these enzymes. Second, K_m_ and K_cat_ of CynD_PER-URP-08_ were 1.93 mM and 6.65 s^−1^, respectively. Third, despite that CynD_PER-URP-08_ and CynD_C1_ only differ in five positions, CynD_PER-URP-08_ retain almost the same activity in pH 9 whereas CynD_C1_ has been reported as almost inactive at this pH. Fourth, as pH is known to influence the oligomerization of CynDs, we reported that in pH 8, CynD_PER-URP-08_ forms spirals made up of an estimated ~24 to ~48 subunits showing that several oligomeric states are present in this pH. This is different compared with CynD_C1_ that was reported to forms just oligomers of 18 subunits at this pH. Moreover, at pH 11, the CynD_PER-URP-08_ monomer was observed. Finally, we showed for the first time, that the abundance of CynD_PER-URP-08_ transcripts increases 6-fold when bacterial cultures are exposed to CN^−^.

Altogether, the results we reported here warrant further investigation to explore the potential of *B. safensis* PER-URP-08 and CynD_PER-URP-08_ for cyanide bioremediation.

## DATA AVAILABILITY

The final genomes assemblies are available in IMG/M (Chen et al., 2021) and GenBank/NCBI (Benson et al., 2013) databases under the accessions numbers: 2818991268, 2818991267, 2818991266 and RSEW00000000.1, RSEY00000000.1, RSEX00000000.1, respectively for *Bacillus safensis* PER-URP-08, *Bacillus licheniformis* PER-URP-12, *Bacillus subtilis* PER-URP-17.

## Acknowledgements

We are very grateful to the Ricardo Palma University High-Performance Computational Cluster (URPHPC) managers Gustavo Adolfo Abarca Valdiviezo and Roxana Paola Mier Hermoza at the Ricardo Palma Informatic Department (OFICIC) for their contribution in programs and remote use configuration of URPHPC. We also thank Germán Sgro for help with negative staining.

## Authors Contributions

Conceptualization: S.J.A., A.G.S., A.M.D.S., Methodology: S.J.A., D.Z.S., A.C.P., M.B.R., C. M.P., L.F.M., P.M.P., C.M. Computing resources: M.Q.A., J.C.S., Data curation: S.J.A., D. Z.S., A.C.P., C.M.P. Formal analysis: S.J.A., D.S.Z., A.C.P., C.M.P., C.S.F. Visualization: S.J.A., D.Z.S. Writing – original draft preparation: S.J.A., Writing – review and editing: S.J.A., J.C.S., C.S.F., A.M.D.S. Supervision: S.J.A., A.G.S., M.Q.A., C.S.F., A.M.D.S. Funding acquisition: A.G.S., M.Q.A., C.S.F., J.C.S., A.M.D.S. All authors read, provided critical review, and approved the final manuscript.

## SUPPLEMENTARY FIGURES

**Figure S1. Proteins containing CN_hydrolase domain in the three genomes studied.** Four CN_hydrolases containing-proteins were identified in the analyzed genomes. YkrU and CynD are present only in *B. safensis* PER-URP-08. MtnU was found in *B. licheniformis* PER-URP-12 and *Bacillus subtilis* PER-URP-17. YhcX was found in the three genomes.

**Figure S2. Alignments of identical protein group CynDs with CynD_PER-URP-08_ and CynD_C1_.** Homology od CynD from *Bacillus safensis* PER-URP-08 is clearly showed in the protein sequence alignments of several CynD homologs including those with tested enzymatic activity as CynD from *B. pumilus* strain C1.

**Figure S3. CynD_PER-URP-08_ production of NH_4_ by time.** Linear adjust of the product formation (NH_4_) by CynD in the first 40 seconds of reaction using different initial concentrations of cyanide.

## SUPLEMENTARY TABLES

**Table S1. Accession numbers of reference genomes used in the assembly process.**

**Table S2. Cyanide removal percentage of the twenty isolates from mine tailings in Peru.**

**Table S3. BLAST best-hits of the partial 16S rRNA gene for each tested strain.**

**Table S4. Summary of IMG/M annotations of the three *Bacillus* genomes.**

**Table S5. Summary information of the 132 genomes used in the core genomes analysis.**

**Table S6. Rhodanese domain coding ORFs in the three sequenced genomes.**

**Table S7. Identical protein groups (IPG) NCBI accession IDs by strain and species.**

