## Supplementary figures and images for "Isolation of a *Bacillus safensis* from mine tailings in Peru, genomic characterization and characterization of its cyanide-degrading enzyme CynD"

### Supplemental_Figure_1

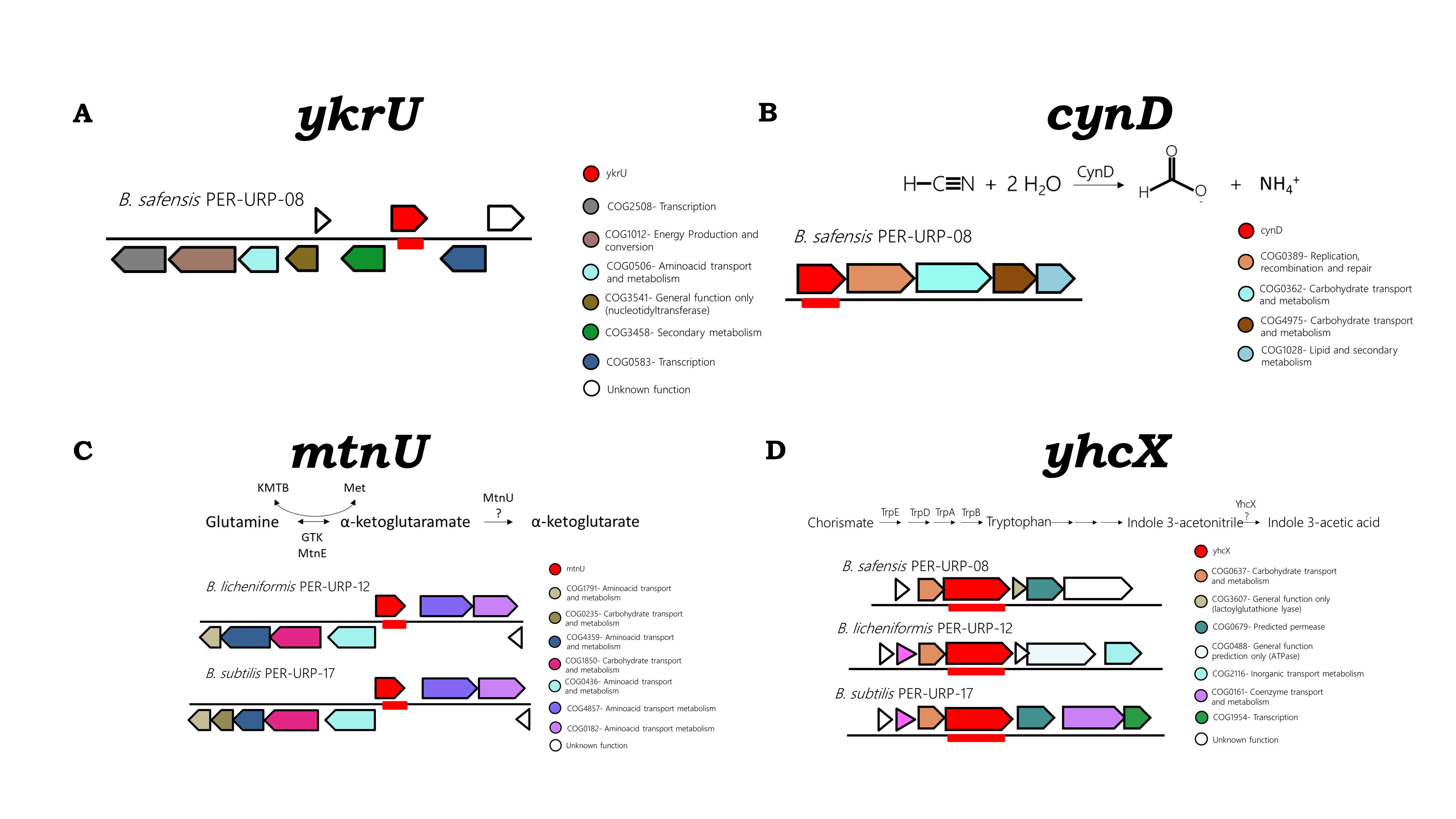

### Supplemental_Figure_3

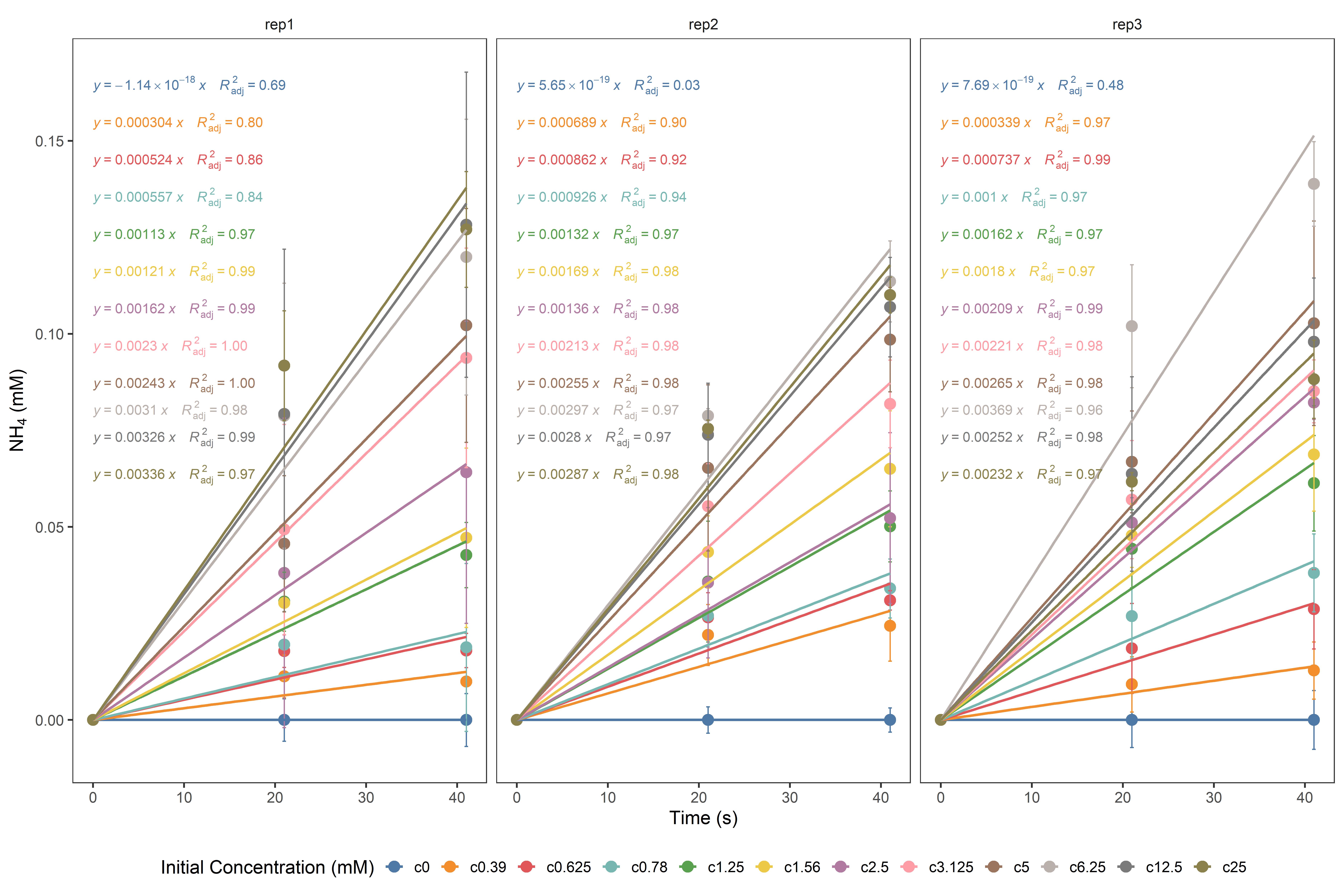
